# Dopamine neurons govern olfactory-gated infant begging behavior

**DOI:** 10.1101/2023.03.18.533277

**Authors:** Julie M. Butler, Devraj Singh, Penelope Baker, Scott V. Edwards, Kyle Summers, Lauren A. O’Connell

## Abstract

Altricial young of many species beg parents for the nutrients required for healthy development. From human crying to begging chicks, young expend precious energy reserves to communicate their hunger. Despite repeated independent evolutionary origins, the neural basis of parent-directed communication by infants is unknown. Here, we examined the sensory and neural basis of begging behavior in tadpoles of the monogamous and biparental Mimetic poison frog (*Ranitomeya imitator*). In this species, tadpoles beg parents for egg meals by dancing. We used this robust motor display to determine that tadpoles use multimodal cues for caregiver recognition, where olfactory cues are necessary for caregiver recognition while visual cues provide an orientation goal. We found that dopamine related brain regions have higher neural activity in begging tadpoles and that dopamine signaling had opposing modulatory effects through D1 and D2 family receptors, similar to swimming behavior in other tadpole species. We then identified caudal posterior tuberculum dopamine neurons as more active during begging behavior and sensitive to caregiver olfactory cues. Projections of these dopaminergic neurons to the spinal accessory motor nucleus are required for begging displays. These findings support the idea that dopamine regulates olfactory-guided parental recognition in young and opens many avenues for studying how new communication behaviors can evolve from ancestral motor circuits.

## Introduction

Communication of nutritional need during infancy is important for acquiring nutrition for healthy development and is among the first social interactions in life. Infants of altricial species often solicit food from their parents, whether this is approaching lactating mammalian mothers for milk [1] or begging for food from provisioning parents in birds and other taxa [2–4]. Across many independent evolutionary origins, begging is an energetically costly motor performance that selects for honest signaling in offspring [5,6]. Due to costs of communication, begging is usually paired with parental recognition, where infants learn to recognize their caregivers or nest sites [7,8]. For example, human infants learn to recognize their mothers apart from other individuals within the first week of life [9,10]. Understanding the neuronal mechanisms controlling begging displays should help elucidate how this essential infant behavior is regulated.

Insights into parent-offspring communication come mostly from studies in mammals and birds [11–13]. While we understand a great deal about this process from the parental perspective [14,15], the infant perspective is unclear, in part because studying infant brains currently comes with many technical challenges. Here, we investigate the neural mechanisms of caregiver-directed communication by tadpoles of the Mimetic poison frog (*Ranitomeya imitator*). In this monogamous and biparental amphibian, parents move individual tadpoles from the leaf litter to pools of water contained within plants [16,17]. In these low-resource nurseries, parents coordinate care by monitoring tadpole nutritional needs, where tadpoles perform a rapid motor display to solicit trophic unfertilized egg meals from their mother [18,19]. Investigating the neural mechanisms of tadpole begging behavior provides an inroad into understanding parent-offspring relationships from the neonate perspective and more broadly how new communication behaviors evolve from existing neural components.

## Results

### Tadpoles recognize caregivers using olfaction

We tested which caregiver-related stimuli elicit begging behaviors (**Fig. 1a**) by exposing tadpoles to control or caregiver visual, chemosensory, or multimodal stimuli. Tadpoles exposed to caregiver multimodal cues had significantly higher begging (*X*^*2*^=46.08, P<0.001; **Fig 1b**) compared to other groups. When presented with mixed multimodal stimuli (caregiver chemosensory stimuli + visual control), tadpoles were also more likely to beg (*X*^*2*^=33.538, P<0.001), and the time spent begging did not differ from those presented matched caregiver multimodal stimuli (F=0.848; P=0.256; **Fig. 1c;** Sup Fig. 1). These patterns suggest olfactory cues are sufficient to elicit begging to a visual object, but the visual object is not required to be a frog.

**Figure 1.**
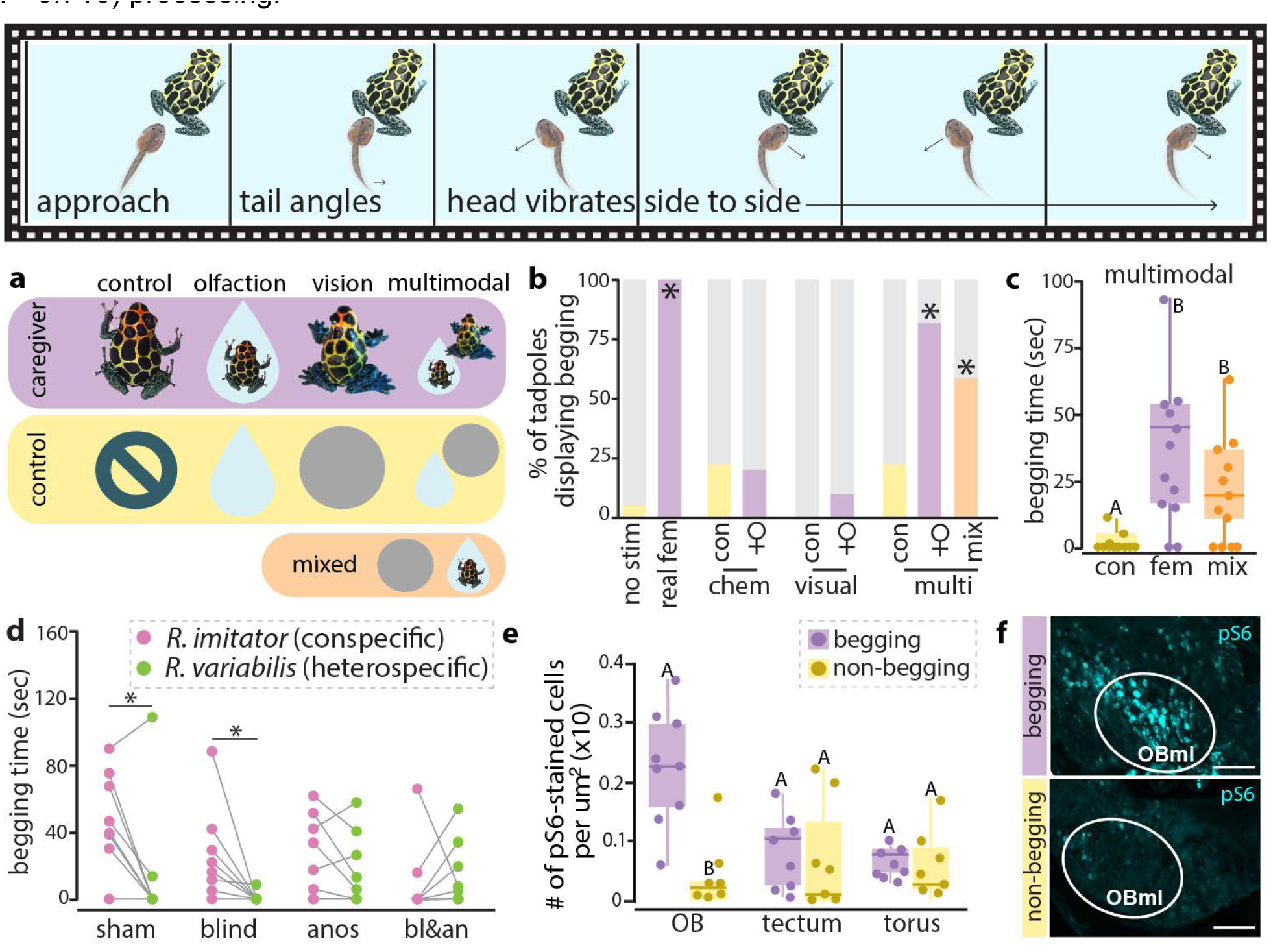
Tadpoles use chemosensory sensory stimuli for caregiver recognition and begging. Top panel shows a schematic time lapse of characteristic begging behavior where tadpoles orient to a caregiver and vibrate their body back and forth. **(a)** Different cue types were presented to tadpoles to determine which sensory modalities are sufficient to elicit begging behavior. **(b-c)** Only tadpoles exposed to a real female, caregiver multimodal stimuli, and mixed multimodal stimuli are significantly more likely to beg compared to the no stimulus group. **(d)** Tadpoles sensory systems were ablated and tadpoles were presented with a female conspecific (*R. imitator*) or a female heterospecific (*R. variabilis*). Sham and blind tadpoles show increased begging towards *R. imitator* females whereas anosmic and blind + anosmic tadpoles do not distinguish between these species. **(e-f)** Begging tadpoles have higher neural activation in the olfactory bulbs than non-begging tadpoles. * indicates significantly higher/lower likelihood to beg compared to control groups (chi-square). Different letters indicate statistical differences. Abbreviations: anos, anosmic; bl&an, blind and anosmic; chem, chemosensory; fem, female; multi, multimodal; OB, olfactory bulbs; OBml, mitral layer of the olfactory bulbs. Scale bars in f represent 50 μm.

We next examined if olfactory or visual cues are necessary for caregiver recognition and begging behavior by generating blind, anosmic, and blind+anosmic tadpoles. Control and blind tadpoles beg for longer (sham: P=0.004; blind: P=0.019) to a caregiver than heterospecific female (**Fig. 1d**), but tadpoles lacking smell beg for similar amounts of time (anosmic: P=0.133; blind+anosmic: P=0.414). This suggests that olfaction is key for sensory discrimination of female stimuli.

To relate sensory stimuli to neuronal signaling, we quantified pS6-positive cells as a proxy of neural activity in sensory processing regions of the brain (**Fig. 1e-f**). Begging tadpoles had more pS6-positive cells in the mitral layer of the olfactory bulbs (OBml) compared to non-begging tadpoles (P<0.001). However, there were no differences in activation between begging and non-begging tadpoles in regions associated with visual (tectum; P=0.643) or acoustic (torus; P=0.716) processing.

### Dopamine signaling stimulates begging behavior

To identify where in the brain begging behavior may be regulated, we quantified pS6-positive cells as a proxy of neural activation in several socially-relevant brain regions of begging and non-begging tadpoles. In general, begging tadpoles had overall more pS6-positive cells than non-begging tadpoles (F=14.977; P<0.001; **Fig. 2a;** Sup. Fig. 2) and there is an interaction between behavior and brain region (F=7.245; P=0.003). Begging tadpoles have more pS6-positive cells in the lateral septum (P=0.003; **Fig. 2c**), medium pallium (P=0.029), striatum (P=0.017), nucleus accumbens (P<0.001), and posterior tuberculum (P<0.001). Begging behavior significantly positively correlates with the number of pS6-positive cells in the lateral septum (R=0.705; P=0.048; **Fig. 2c**) and posterior tuberculum (R=0.845; P=0.008).

**Figure 2.**
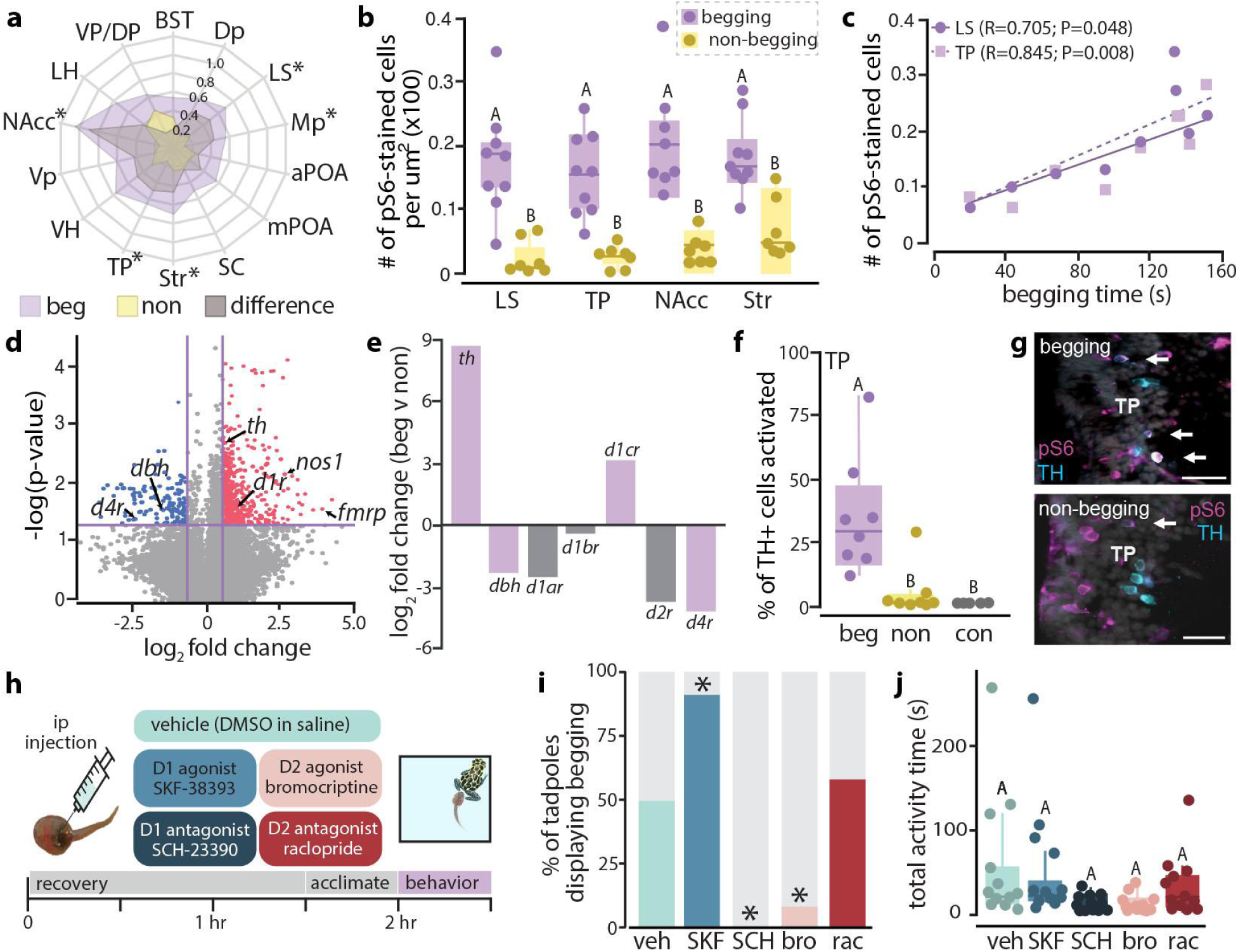
Dopamine mediates tadpole begging behaviors. **(a)** Begging tadpoles have higher neural activation overall, where the further out the net extends, the higher the normalized p-S6 positive cells. Difference is the non-begging average subtracted from the begging average. **(b)** Begging tadpoles have significantly more pS6-positive cells compared to non-begging tadpoles in the lateral septum (LS), posterior tuberculum (TP), nucleus accumbens (NAcc), striatum (Str), and medial pallium (MP). **(c)** Neural activity in the LS (circles) and TP (squares) significantly positively correlates with time spent begging. **(d-e)**. PhosphoTRAP identified the dopamine system as a potential mediator of begging, as tyrosine hydroxylase (TH), dopamine beta hydroxylase (DBH), and dopamine receptors are differentially expressed between begging and non-begging tadpoles. Dots represent depleted (blue) or enriched (red) transcripts in begging tadpoles by comparing immunoprecipitated mRNA to total input. Purple bars indicate statistically different expression between begging and non-begging tadpoles. **(f-g)** Begging tadpoles have higher activation of dopaminergic (TH-positive) neurons in the TP compared to non-begging and control tadpoles. **(h-j)**. Tadpoles injected with the D1 agonist SKF are more likely to beg, but tadpoles injected with a D1 antagonist never beg. D2 agonism also inhibits begging behavior. While manipulating dopamine signaling impacts begging, it does not significantly affect overall activity levels. * indicates significantly higher/lower likelihood to beg compared to control groups (chi-square). Different letters indicate statistical differences. Abbreviations: aPOA, anterior preoptic area; BST, basolateral nucleus of the stria terminalis; bro, bromocriptine; con, control; *d1ar*-*d1ac*, dopamine receptor 1 type a, b, and c; *d2r*, dopamine receptor 2; *d4r*; dopamine receptor 4; *dbh*, dopamine beta hydroxylase; Dp, dorsal pallium; LH, lateral hypothalamus; LS, lateral septum; Mp, medial pallium; mPOA, medial preoptic area; NAcc, nucleus accumbens; non, non-begging; *nos1*, nitric oxide synthase 1; rac, raclopride; SC, suprachiasmatic nucleus; Str, striatum; TP, posterior tuberculum; veh, vehicle; VH, ventral hypothalamus; Vp, ventral pallium; VP/DP, ventral and dorsal pallidum.

To probe the molecular signatures of begging behavior in the brain, we used phosphoTRAP on whole brains from begging and non-begging tadpoles (**Fig. 2d;** Sup. Fig. 3). Within begging tadpoles, enriched transcripts included nitric oxide synthase, the fragile X protein, and several genes associated with the dopamine system. Specifically, tyrosine hydroxylase (TH), the rate limiting enzyme in dopamine synthesis, and dopamine receptor D1 (*d1cr*) were enriched in begging tadpoles, with TH showing a 9-fold enrichment compared to non-begging tadpoles (**Fig. 2e**). Dopamine beta hydroxylase (DBH), which converts dopamine into norepinephrine, and D2-family receptors were depleted in begging tadpoles. Together, this suggests increased dopamine signaling in begging behavior through D1-family receptors. We then confirmed that begging tadpoles had more TH-positive neurons that colocalized with pS6 in the posterior tuberculum and preoptic area (beg v non: F_2,38_=21.643, P<0.001; region: F_2,38_=0.1437, P=0.250; **Fig. 2f-g**; Sup. Fig. 4) compared to non-begging and control tadpoles.

We next tested if tadpole begging relies on dopaminergic signaling using pharmacology (**Fig. 2h**). Tadpoles injected with a D1R agonist begged significantly more frequently than\ vehicle-injected tadpoles (**Fig. 2i**; *X*^*2*^=5.042, P=0.025), but tadpoles injected with a D1R antagonist never displayed begging. Injecting tadpoles with a D2 agonist also inhibited begging behavior, but the D2R antagonist had no impact (*X*^*2*^=0.168, P=0.682). None of the drugs used affected overall activity levels (F_4,55_=2.510, P=0.053; **Fig. 2j**) or aggressive behavior (F_4,15_=2.280, P=0.109; Sup. Fig. 5).

### Dopaminergic neurons are sensitive to caregiver odors

To investigate if caregiver chemosensory stimuli are sufficient to activate dopaminergic neurons in the brain, we exposed tadpoles to unconditioned water (control) and water conditioned with either reproductive *R. imitator* or *R. variabilis* females. Tadpoles exposed to water from a natural caregiver were more active (F=6.273, P=0.008; **Fig. 3a**) than tadpoles exposed to *R. variabilis* (P=0.012) and control water (P=0.047). Tadpoles exposed to *R. imitator* water also had higher activation of dopamine neurons in the POA (F=5.739, P=0.010; **Fig. 3b-c**) compared to tadpoles exposed to *R. variabilis* water (P=0.011) but not control water (P=0.131). While there were no difference of activation of dopamine neurons in the rostral part of the TP, tadpoles exposed to caregiver-conditioned water had significantly higher dopaminergic neuron activation in the caudal TP (F=18.755, P<0.001) compared to those exposed to control (P<0.001) and *R. variabilis* (P<0.001) water. Together, this suggests dopaminergic neurons in the posterior tuberculum are sensitive to caregiver olfactory cues.

**Figure 3.**
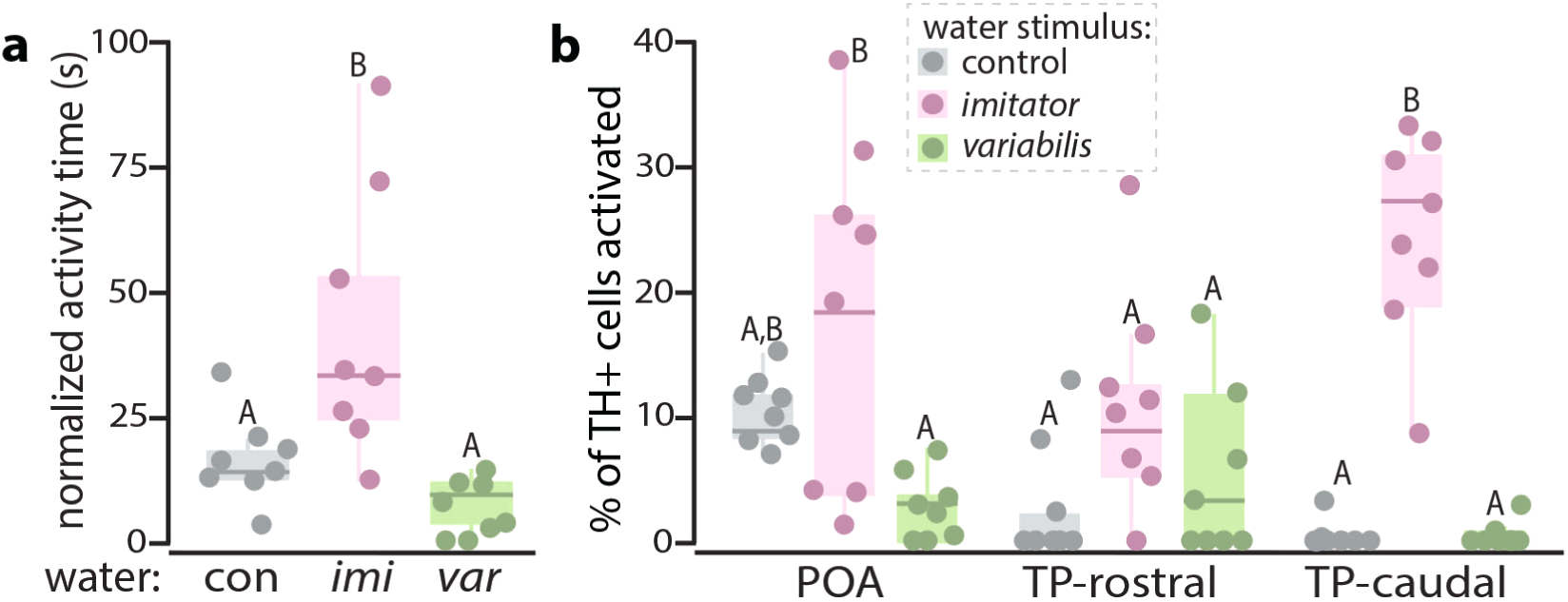
Linking caregiver smells to dopamine neurons. **(a)** Tadpoles exposed to caregiver water (*imi*) are more active than those exposed to control and non-caregiver water (*var*). **(b)** Tadpoles exposed to caregiver smells have higher activation of dopaminergic neurons in the caudal portion of the TP compared to control tadpoles and those exposed to non-caregiver water. Abbreviations: con, control unconditioned water; *imi, R. imitator* conditioned water; POA, preoptic area; TP, posterior tuberculum; *var, R. variabilis* conditioned water.

### Dopaminergic projections to the spinal accessory nucleus are required for begging

Tadpole begging is a motor behavior and we searched for motor neurons in the spinal cord that were near TH-positive fibers and had more pS6-positive cells in begging tadpoles compared to controls. At the junction of the spinal cord and hindbrain, we found more pS6-positive cells in the spinal accessory nucleus (nmXl) in begging tadpoles compared to non-begging and control tadpoles (F_2,19_=10.522, P<0.001; **Fig. 4a**). This location has dense dopamine innervation (Sup. Fig. 6) and controls head and neck movements in mammals [20].

We next functionally tested the role of dopaminergic projections to the nmXl with 6-hydroxydopamine (6-OHDA), a toxin that ablates dopaminergic neurons [21]. Injection of 6-OHDA into the nmXI selectively ablated caudal TP dopaminergic neurons (P<0.001; **Fig. 4b**), but not rostral TP dopaminergic neurons. Tadpoles with ablated cTP dopamine neurons had a significantly reduced chance of begging compared to control tadpoles, with 88% of vehicle-injected tadpoles displaying begging, but only 1 of the 6-OHDA-injected tadpoles displayed a single begging bout for 3 sec (*X*^*2*^=9.000, P=0.027; begging time: P=0.019; **Fig. 4c**). Treatment did not affect overall activity compared to controls (nmXI: P=0.072; **Fig. 4d**).

**Figure 4.**
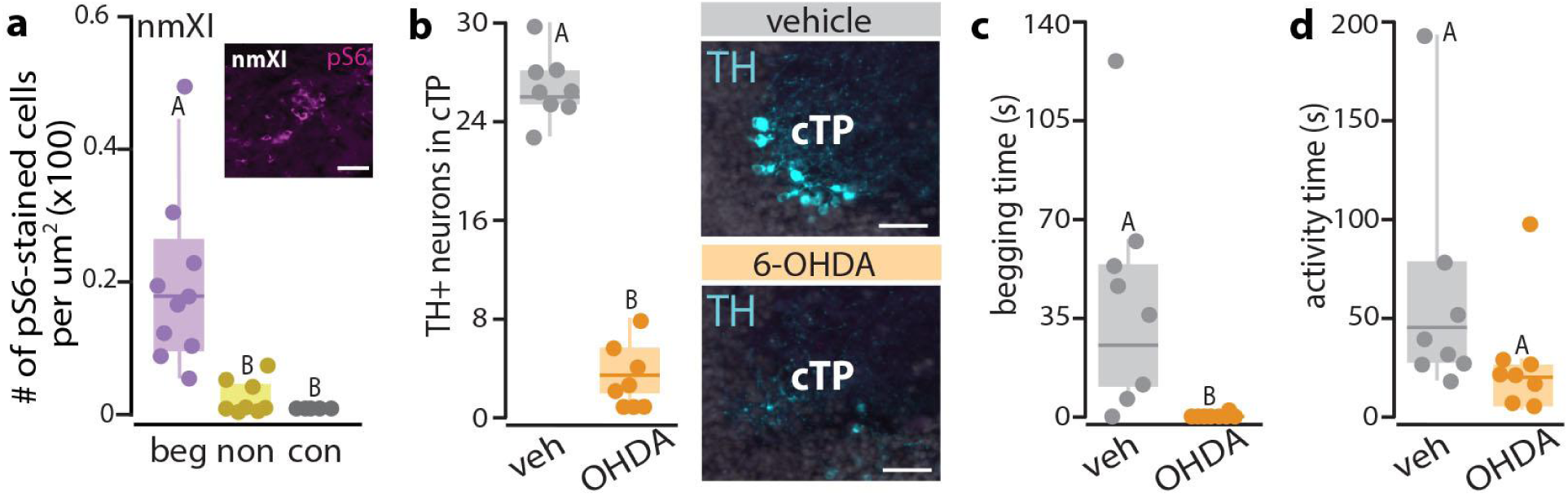
Dopaminergic projections to nmXl are required for begging behavior. **(a)** Activation in the spinal accessory motor nucleus (nmXI) is higher in begging tadpoles compared to control and non-begging tadpoles. **(b)** Injections of the dopaminergic neuron toxin 6-OHDA into the nmXI selectively ablated caudal TP (cTP) dopamine neurons. **(c)** Ablation of cTP dopamine neurons projecting to the nmXI prevented begging behavior compared to controls. **(d)** Overall activity was not impacted by the absence of cTP DA neurons. Different letters indicated statistical differences. Scale bars represent 50 um. Abbreviations: 6-OHDA, 6-hydroxydopamine hydrobromide; beg, begging tadpoles; con, conditional water control; cTP, caudal posterior tuberculum; non, non-begging tadpole; veh, vehicle.

## Discussion

In this study, we have identified the sensory system logic underlying recognition of potential caregivers. We show that recognition of caregivers is mediated by olfaction in *R. imitator* tadpoles, similar to human infants [9,10] and other young mammals [22] that learn the smell of their mothers within the first week of life. We initially hypothesized that recognition of a potential caregiver would rely on vision rather than olfaction based on previous studies in the Strawberry poison frog (*Oophaga pumilio*) showing visual cues are necessary for tadpole begging [23] and influence mate choice later in life [24]. However, newborns and young of many species have limited or absent visual capabilities at birth/hatching [25], including tadpoles [26]. We found tadpoles would beg to a control object if paired with female conspecific odors and blind tadpoles still display begging behavior. However, chemosensory cues alone were not sufficient for begging, suggesting olfaction is important for stimulus identity and visual cues provide an object to direct their begging effort. The need for multimodal stimuli to elicit begging may reflect the high cost associated with the display [19].

Communication of nutritional need is important in altricial young and we provide evidence that dopaminergic signaling is important for begging displays that elicit meals from caregivers.

Dopamine plays many roles in social and motor behavior across taxa [27,28]. In fish and tadpoles, dopamine regulates swimming behavior, with signaling through D1 family receptors promoting swimming and signaling through D2 family receptors inhibiting swimming [29].

Similarly, we found begging behavior was simulated with D1 family agonists and inhibited by D2 family agonists. However, neither total activity as a proxy of swimming behavior or aggression was affected in our dopamine pharmacology experiments. In the wild, *R. imitator* tadpoles are in tiny pools contained within plants and likely perform little swimming behaviors whereas begging is required to elicit food from their parents. It is possible that tadpole begging behavior represents a neofunctionalization of swimming motor circuitry similar to how birdsong evolved from ancestral motor learning circuits [30,31].

We found that dopaminergic projections from the ventral diencephalic dopaminergic neuron populations to the spinal accessory nucleus (nmXl) are required for begging behavior. Most research on dopamine and social motor behaviors are focused on midbrain dopaminergic neurons (A8-10 in mammals) [32]. However, the main source of dopaminergic input to the forebrain and hindbrain in amphibians and fish is the posterior tuberculum, which is homologous to the dorsal hypothalamus and causal thalamus in mammals (A11 group) [33]. In zebrafish, these A11-like neurons are tuned to different sensory stimuli, specifically mechanosensory and visual inputs [34]. Our work provides evidence that these neurons are sensitive to socially-relevant olfactory information, where caudal TP neurons were more active when presented with odors of a potential caregiver compared to a heterospecific female. However, this was not sufficient to stimulate begging, suggesting a “threshold” must be reached, possibly by pairing olfactory stimuli with a visual goal. Ablating the projections of these caudal TP neurons to the nmXl motor nucleus inhibited begging behavior, suggesting this connection is crucial for begging displays. Mouse pups with 6-OHDA dopamine depletions receive less maternal care [35], although it is unclear if this is due to impaired communication.

Overall, tadpole begging behavior opens up numerous opportunities to explore the complex interactions between parents and offspring and supports the idea that new communication behaviors can evolve from ancestral motor circuitry.

## Methods

### Animals

*Ranitomeya imitator* tadpoles were captive bred in our poison frog colony from adult breeding pairs. Tadpoles were approximately Gosner stage 26 (hindlimb development less than 0.5mm) at the time of behavior trials. We used adult *R. imitator* or *R. variabilis* females from actively reproducing breeding as conspecific and heterospecific stimulus animals respectively. All procedures were approved by Harvard University Animal Care and Use Committee (protocol #17-02-293) and Stanford Administrative Panel on Laboratory Animal Care (protocol #33097).

### Begging behavior assays

To examine begging behaviors, we exposed naive tadpoles to reproductive females as described previously [26]. After an acclimation period, tadpoles were exposed to a reproductive *R. imitator* female and allowed them to interact for 20 minutes. Behaviors were recorded from above using a GoPro camera and were later scored by an observer unaware of tadpole identity using BORIS [36] to quantify swimming, begging, and proximity to the female. At the conclusion of the trial, the female was quickly removed from the arena, the light turned off, and the tadpole incubated in the arena for 30 min. We collected and processed brains as previously described for immunohistochemistry and phosphoTRAP [26].

### PhosphoTRAP

We used phosphoTRAP to molecularly profile active neurons in the brain during begging. All samples were processed and sequenced as previously described [26,37,38]. Brains from three tadpoles were pooled together for each sample for phosphoTRAP. Sequencing results were aligned to a *R. imitator* transcriptome, and count data was analyzed in R using paired t-tests on each transcript between the input and immunoprecipitated samples. Fold changes were also calculated for each transcript as log2 (IP count / TOT count). Differentially expressed genes were defined as having a p-value under 0.05 and a fold change greater than 1.5 in either direction.

### Sensory system assays

We exposed tadpoles to a caregiver visual (frog model) and chemosensory (female-incubated water) stimuli. Controls included a gray disk (visual) or plain frog water. Additional controls included a negative control (no stimulus) and positive control (live reproductive *R. imitator* female). Behavior assays were conducted as above except that we recorded a 10 minute baseline for each tadpole and then presented it with a randomly assigned unimodal visual or chemical stimulus for 20 min. We then added a second unimodal stimulus or control of the other modality to create a multimodal experience. In all, there were 9 different exposure groups: negative control, positive control (real female), control visual, caregiver visual, control chemosensory, caregiver chemosensory, multimodal control, multimodal caregiver, multimodal mix (caregiver chemosensory with visual control). At least 10 tadpoles were exposed to each stimulus, with all tadpoles receiving two different stimuli conditions. Order of the stimulus was randomized and did not appear to influence behavior.

We also collected a group of tadpoles exposed to control and conditioned water in the absence of other stimuli. *Ranitomeya imitator* and *R. variabilis*-conditioned water were generated by incubating reproductive females in frog water for 4 hours. Tadpoles were tested for begging behavior after a 90 min acclimation followed by exposure to a randomly assigned conditioned water for 40 min.

### Sensory ablations

We generated blind tadpoles by removing eyes and anosmic tadpoles by severing their olfactory nerves just caudal to the olfactory epithelium. Double-ablated tadpoles underwent both surgeries, and sham-control tadpoles were anesthetized and received a small incision on each flank. All tadpoles were allowed to recover for 1 week before use in behavior trials and displayed normal swimming and eating behaviors. Tadpoles were tested in a begging assay as described above except using a live reproductive *R. imitator* or *R. variabilis* female in random order with a 20-min recovery and water change between the two exposures.

### Immunohistochemistry

To compare neural activation in the brains of begging and non-begging tadpoles, we stained cryosectioned tadpole brains for phosphorylated ribosomes [38] (pS6, a proxy for neural activity) as previously described [26]. We used co-stained for pS6 and tyrosine hydroxylase (TH, the rate-limiting step in dopamine synthesis) with mixtures of primary antibodies [1:5000 rabbit anti-pS6 (Invitrogen; cat# 44-923G); 1:500 mouse anti-tyrosine hydroxylase (EMD Millipore; cat# MAB318)] and secondary antibodies (1:250 AlexaFlour 488 anti-mouse and 1:250 AlexaFlour 647 anti-rabbit).

Stained brain sections were imaged on a Leica DM6B microscope, and the number of labeled cells was quantified in FIJI image analysis software [39] as previously described [26]. We identified brain regions based on cytoarchitectural landmarks, and comparisons with a poison frog brain atlas [37]. pS6-positive cells were quantified in 14 socially-relevant brain regions, 4 sensory processing regions, and one motor region. Colocalization of TH and pS6-positive cells were quantified in the preoptic area and posterior tuberculum.

### Dopamine Pharmacology

Tadpoles were anesthetized in buffered 0.01% MS-222 in frog water and given an intraperitoneal (ip) injection of a selective D1 family agonist (10ug/gbm; SKF 38393; Sigma, #D047) or antagonist (0.5ug/gbm; SCH-23390 hydrochloride; Sigma, #5057230001), D2 family agonist (20ug/gbm; bromocriptine; Sigma #B2134) or antagonist (5ug/gbm; raclopride; Sigma, #5058770001), or vehicle (DMSO; Sigma, #D8418) diluted in 0.9% sterile saline solution.

Tadpoles were allowed to recover for 2 hours before begging behavior trials were run as described above. All tadpoles were awake and swimming around their arena within 30 min of the injection. As an additional control, we subjected a subset of tadpoles (4 per group) to a smaller tadpole (Gosner stage 25) for 10 min and quantified the number of aggressive behaviors displayed in addition to affiliative behaviors.

### 6-OHDA manipulations

We used the dopamine toxin 6-hydroxydopamine-hydrobromide (Sigma, #162957) to ablate dopamine neurons in the hindbrain and striatum by injecting anesthetized tadpoles with ∼50 nl of OHDA or saline into target regions of each hemisphere using a Nanoject. Tadpoles were placed back into their home containers and allowed to recover. All animals were tested 7 days post-injection in begging behavior assays and euthanized for tissue processing as described above. At the developmental stage used for these experiments (Gosner stages 26-30), noradrenergic neurons are not reliably detectable via immunostaining for dopamine beta hydroxylase. However, TH-immunoreactive neurons were occasionally observed in the isthmal region and near the solitary tract in the caudal hindbrain of both control and OHDA tadpoles with no apparent differences between the groups, suggesting the OHDA injections did not impact noradrenergic neurons.

### Statistical Analysis

All statistics were performed in R (v4.0.2). We used student’s t-tests and generalized linear mixed models (GLMM; Package: glmmTMB; [40]) to compare behaviors among groups, with animal ID as a repeated, random factor when appropriate. Cell count data was analyzed using a GLMM with brain region as a repeated factor and animal ID as a random factor. To test the likelihood of begging, we use chi square tests to test each group against the appropriate experimental control group. When appropriate, Tukey’s post-hoc tests were used to parse differences among groups. Correlations were done using Pearson correlations. Statistical outliers were detected and removed using an Iglewicz and Hoaglin’s test with a modified z-score of 3.5 prior to analyses. All graphs were produced in R using ggplot2 [41] and assembled into figures using Adobe Illustrator (version 2023).

## Acknowledgements

We thank members of the O’Connell lab for assistance with animal care and Liqun Luo discussions about this research.

