## Supplementary Figures for "Dopamine neurons govern olfactory-gated infant begging behavior"

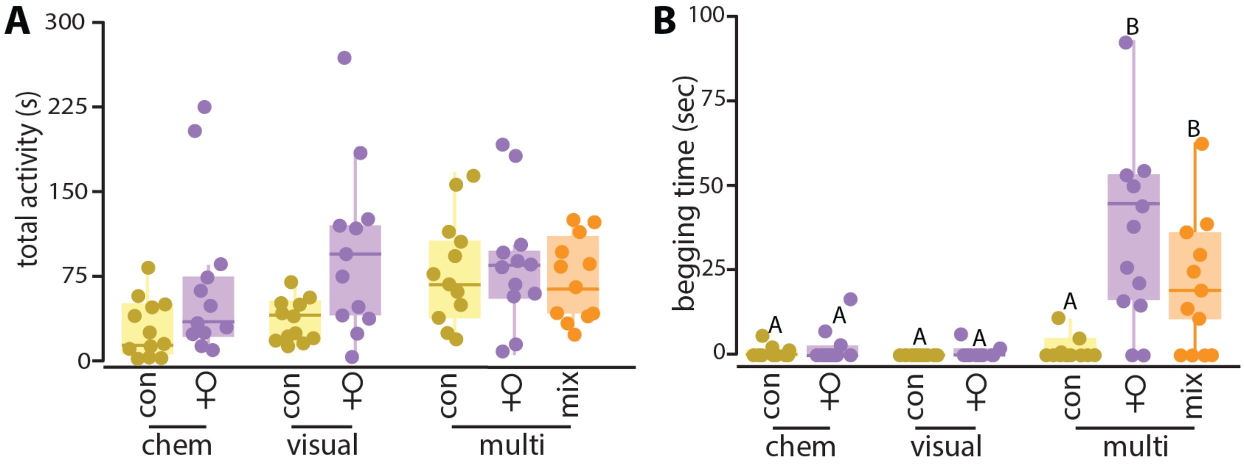


**Supplemental Figure 1. Multimodal caregiver stimuli increase begging behavior.** Tadpoles exposed to caregiver and mixed multimodal stimuli beg for more time than tadpoles exposed to unimodal stimuli or control multimodal stimuli, but there is no impact on total activity.

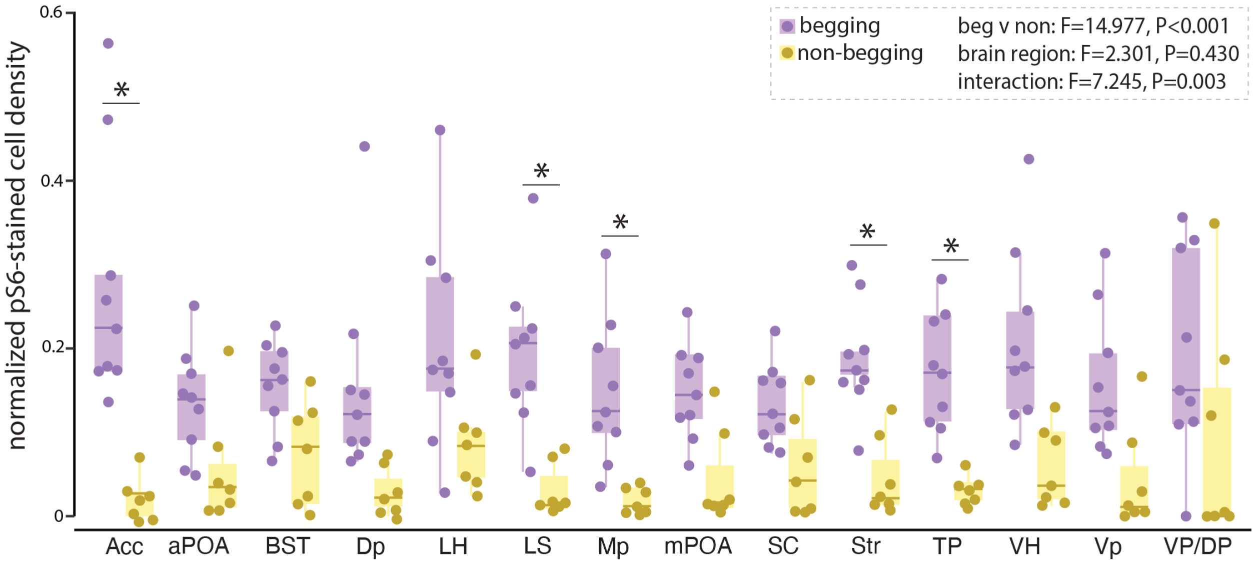


**Supplemental Figure 2. pS6 staining is generally higher in begging tadpoles compared to non-begging tadpoles.** * represent statistical differences within a region between begging and non-begging tadpoles. Abbreviations: aPOA, anterior preoptic area; BST, basolateral nucleus of the stria terminalis; Dp, dorsal pallium; LH, lateral hypothalamus; LS, lateral septum; Mp, medial pallium; mPOA, medial preoptic area; NAcc, nucleus accumbens; SC, suprachiasmatic nucleus; Str, striatum; TP, posterior tuberculum; VH, ventral hypothalamus; Vp, ventral pallium; VP/DP, ventral and dorsal pallidum.

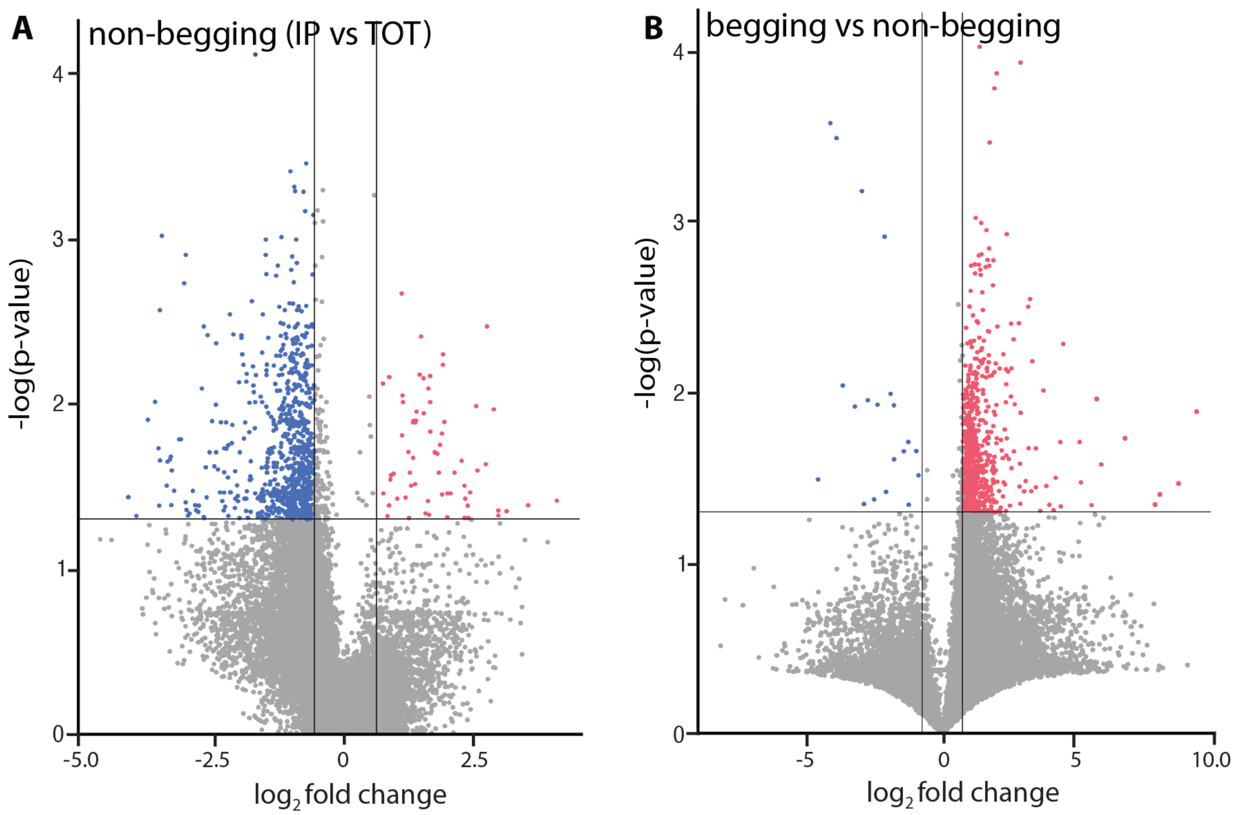


**Supplemental Figure 3. Volcano plots from phosphoTRAP analyses. A.** Non-begging tadpoles have a higher number of depleted transcripts (blue) compared to enhanced transcripts (red). **B**. We also compared IP/TOT ratios between begging and non-begging tadpoles, which yields 503 transcripts up-regulated in begging tadpoles compared to non-begging tadpoles, and 22 transcripts down-regulated in begging compared to non-begging tadpoles.

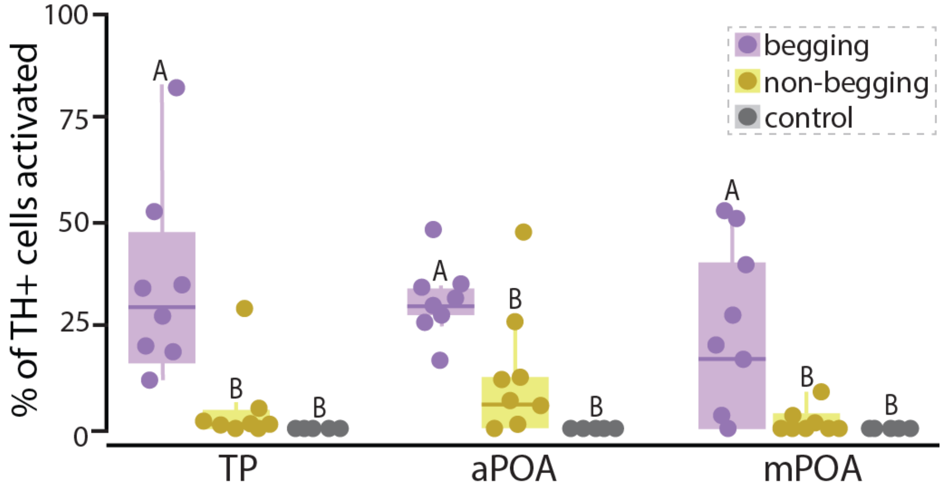


**Supplemental Figure 4. Begging tadpoles have higher activation of dopamine neurons in the TP, a POA and mPOA compared to non-begging and control tadpoles.** Different letters indicated statistical differences.

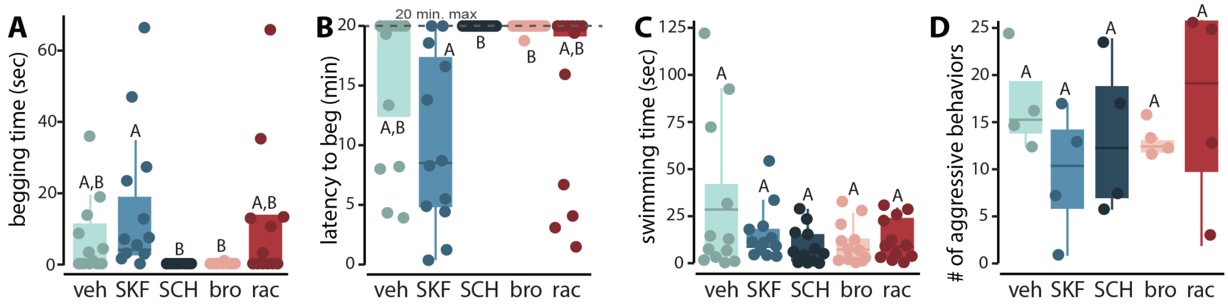


**Supplemental Figure 5. D1R signaling modulates begging behavior. A-B.** Tadpoles injected with the D1R agonist SKF beg more and sooner compared to tadpoles injected with the D1R antagonist SCH. The D2R agonist bromocriptine also reduced begging behavior compared to the D1R agonist tadpoles. C-D. None of the agonists or antagonists used affected swimming behavior or the number of aggressive behaviors directed towards a smaller conspecific. Different letters indicate statistical significance. Abbreviations: bro, bromocriptine; rac, raclopride (D2R antagonist); SCH, SCH-23390; SKF, SKF-38393; veh, vehicle.

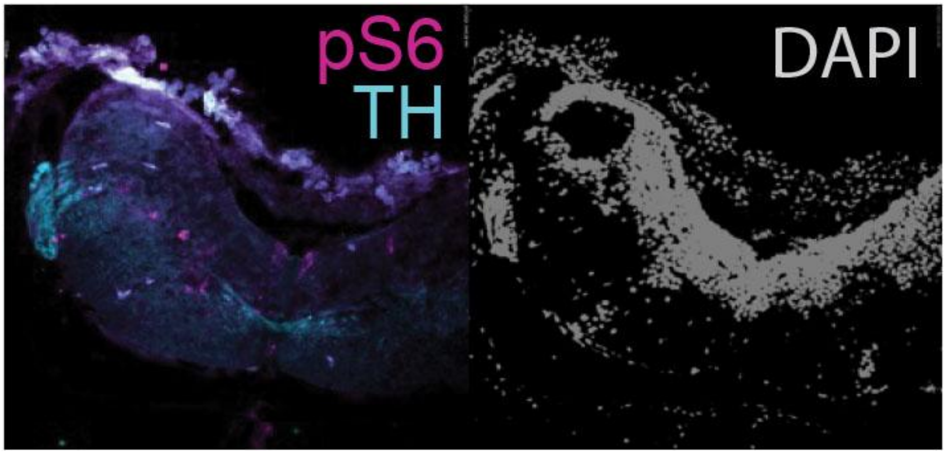


**Supplemental Figure 6. The spinal accessory nucleus (nmXI) has tyrosine hydroxylase-positive fibers.**
